# On the emergence of *Candida auris*: climate change, azoles, swamps and birds

**DOI:** 10.1101/657635

**Authors:** Arturo Casadevall, Dimitrios P. Kontoyiannis, Vincent Robert

## Abstract

The most enigmatic aspect of the rise of *Candida auris* as a human pathogen is that it emerged simultaneously in three continents with each clade being genetically distinct. Although new pathogenic fungal species are described regularly, these are mostly species associated with single cases in individuals who are immunosuppressed. In this study, we used phylogenetic analysis to compare C. *auris* with temperature susceptibility of close relatives and use these results to argue that it may be the first example of a new fungal disease emerging from climate change with the caveat that many other factors could have contributed.

---

*Candida auris* is a new drug resistant fungal species that was first isolated in 2009 from a human ear and thus named ‘auris’ (1). Since then, *C. auris* has been associated with human disease in many countries and the clinical isolates are remarkable for exhibiting non-susceptibility to antifungal agents. One of the striking developments associated with the appearance of *C. auris*-related disease is that pathogenic isolates appear to have emerged independently in three continents simultaneously (2). Analysis of isolates recovered from the Indian subcontinent, Venezuela and South Africa during 2012-2015 revealed that the isolates from each continent were clonal but those from different continents constituted genetically different clades (2). The mechanism(s) responsible for the simultaneous emergence of three different clades of *C. auris* in three geographically distant regions are unexplained.

The use of widespread use of antifungal drugs has been suggested as a contributory cause in the emergence of *C. auris* (3). Whereas selection by environmental azole use can certainly have contributed to drug resistance in this fungal species, it does not easily explain why this organism suddenly became a human pathogen in three continents. For example, the emergence of azole resistant *Candida* spp. began long before the appearance of *C. auris* and there does not appear to be a correlation between the emergence of azole resistant *Aspergillus* spp. associated with azole agricultural use and the hot spots for *C. auris* emergence. The acquisition of drug resistance alone is very unlikely to confer upon a microbe the capacity for pathogenicity, since reduced susceptibility to drugs and virulence are very different properties as evidenced by frequent fitness cost associated with mutations conferring resistance of *Candida* to antifungals (4,5). Instead, as exemplified by the experience with *Aspergillus fumigatus*, aspergillosis was a well-known clinical entity before the putative acquisition of drug resistance from agricultural use of azoles (6). Hence, one might have expected that *C. auris* to have been known as a human and animal pathogen first, and then acquired drug resistance, rather than simultaneously emerging as a drug resistant human pathogen from agricultural drug use. Another suggested explanation for the emergence of *C. auris* is that it recently acquired virulence traits that conferred the capacity for virulence (7). Although this explanation cannot be ruled out, the capacity for virulence is a complex property that emerges from many attributes, and it is improbable that it would have occurred concurrently in three continents, unless driven by another factor that selected for it.

Human pathogenic fungi constitute only a very small minority of an enormous number of fungal species in the environment, numbering in the few hundred (8). In contrast to ectothermic animals and plants, mammals are remarkably resistant to invasive fungal diseases. Mammalian resistance to invasive fungal diseases is proposed to result from a combination of high basal temperatures that create a thermal restriction zone and advanced host defense mechanisms in the form of adaptive and innate immunity (9). In this regard the emergence of such thermodynamically unfavorable animals such as mammals as the dominant large animals has been proposed to have resulted from a fungal filter at the Cretaceous-Tertiary boundary that prevented a second reptilian age (9). According to this theory, fungal pathogens are rare in mammals because this group of animals was selected by the fungi at the end of the Cretaceous (9). Supporting this view is the fact that the majority of fungi grow well in ambient temperatures but only a small percentage of species can replicate at 37 °C (10). Consequently, invasive fungal infections are rare unless one of these two resistance pillars is disturbed. For example, the high prevalence of mycoses in individuals with advanced HIV infection was a result of weakening of the immune system, while the white nose syndrome in bats occurs during their hibernation process when their temperature drops (11).

The thermal restriction zone that protects mammals is the difference between their high basal temperatures and the environmental temperatures. Human induced climate change is anticipated to warm earth by several degrees in the 21^st^ century, which will reduce the magnitude of the gradient between ambient temperatures and mammalian basal temperatures (12). Consequently, there is concern that higher ambient temperatures will lead to the selection of fungal lineages to become more thermally tolerant such that they can breach the mammalian thermal restriction zone. In this regard, experiments with entomopathogenic fungus have shown that these can be rapidly adapted to growth at higher temperatures by thermal selection (13), establishing a precedent for an animal pathogenic fungus adapting to mammalian basal temperatures. Most worrisome, analysis of thermal tolerances for fungal isolates deposited in a culture collection showed a trend in increased capacity to replicate at higher temperatures for basidiomycetes, consistent with an early adaptation to higher ambient temperatures beginning in the late 20 ^th^ century (14) and fungal species in cities have become more thermo tolerant than their rural counterparts (15). An analysis of the correlation of temperature tolerance with latitude (for strains isolated in recent decades and described in (14)) showed that the Pearson correlation between the maximum temperature growth of all fungi, ascomycetes and basidiomycetes was −0.14619, −0.054638414, and −0.304251976, respectively. The lack of a significant trend for the ascomycetes is consistent with the fact that this group can grow at higher ambient temperatures and can thus grow across the latitudes. For the basidiomycetes, the borderline significant trend suggests that these are adapting to the warmer conditions in higher absolute latitudes, possibly because of climate warming, consistent with our earlier analysis (14). Given the capacity of fungal species to adapt to higher temperatures, and the fact that many fungal species that are currently non-pathogenic species are likely to have the necessary virulence attributes by virtue of their survival in soils, we previously hypothesized that climate change would bring new fungal diseases (12).

*C. auris* is an ascomycetous yeast and a close relative of *Candida haemulonii* species complex, which includes species occasionally pathogenic in humans and animals and demonstrates a high level of baseline antifungal drug resistance (16). This phylogenetic connection could explain its low susceptibility to antifungal agents and the possession of virulence attributes that confer it with pathogenic potential. To evaluate our hypothesis we compared the thermal susceptibility of *C. auris* to some of its close phylogenetic relatives and found that the majority of these were not tolerant for mammalian temperatures (Figure 1). Although this tree reveals that *C. auris* is capable of growing at higher temperatures than most of its closely related species, it does not inform as to whether this is a new trait. It is noteworthy that the earliest description of *C. auris* came from a strain recovered from a human ear, which is much cooler than core body temperatures. Hence, this fungus may have gone through a short transient period where it inhabited human surfaces before being associated with disease. Currently, *C. auris* preferentially colonizes the cooler skin rather than the hotter gut mycobiome, a preference that may be consistent with a recent acquisition of thermo tolerance.

**Figure 1.**
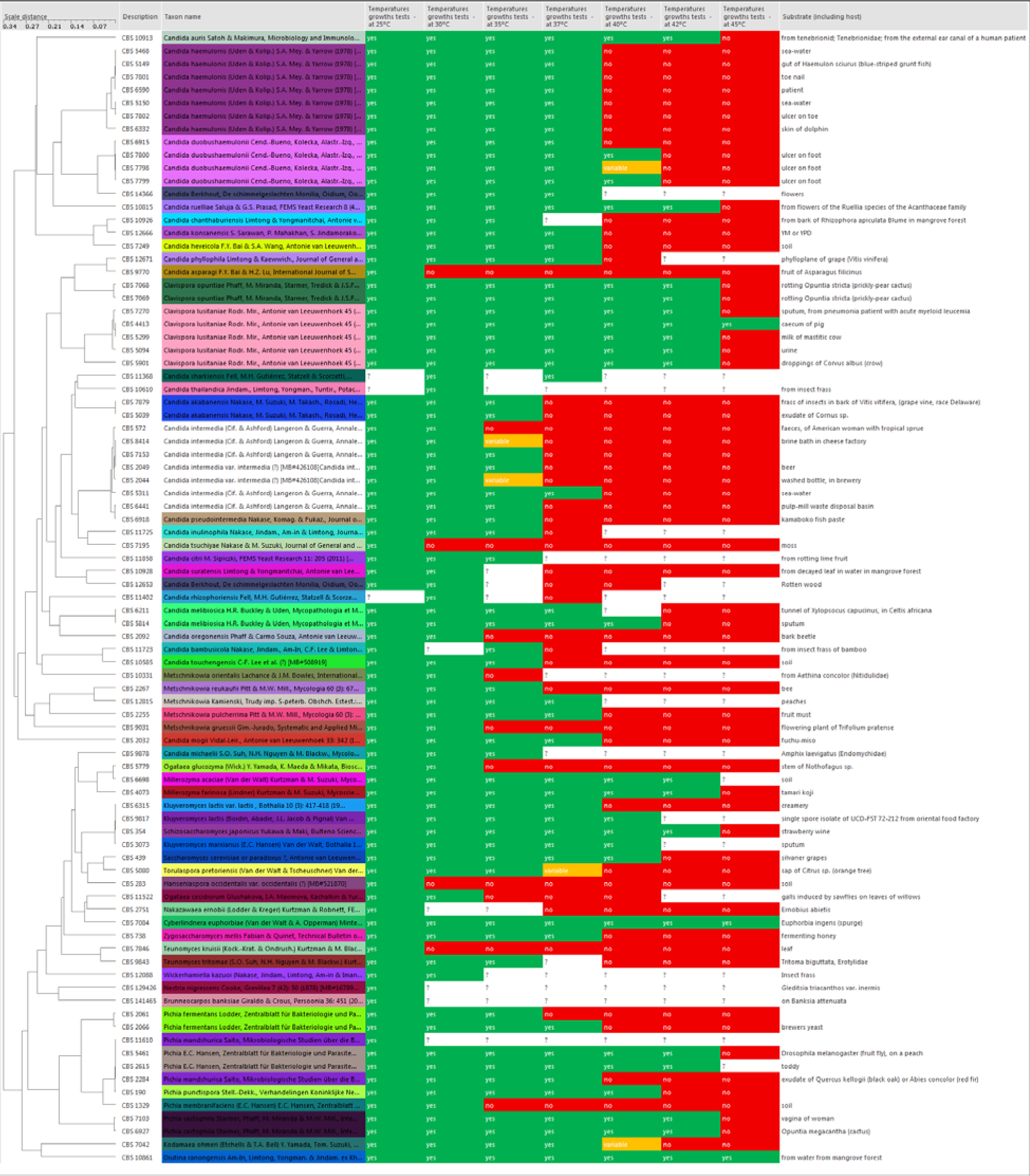
Comparison of thermal tolerance of *C. auris* and several close relatives. The tree shows that thermal tolerance is not monophyletic to closely related species. Most closely related species manifest lower thermal tolerance that *C. auris*. The hierarchical tree (UPGMA) is based on pairwise sequence alignments of both Internal Transcribed Spacers and Large Subunit Ribosomal DNA (data obtained and available from the CBS collection at www.westerdijkinstitute.nl).

With this background, we propose the hypothesis that *Candida auris* is the first example of a new pathogenic fungus emerging from human induced global warming. We posit that prior to its recognition as a human pathogen, *C. auris* was an environmental fungus. The fact that *C. auris* fails to grow anaerobically, along with the fact that is typically detected in cooler skin sites but not the gut, supports the notion that *C. auris* was an environmental fungus, until recently. Several factors, not necessarily mutually exclusive, could have been operative on why *C. auris* emerged in the last decade. For example, as *C. auris* constitutively overexpresses HSP 90, this could account for both its multidrug resistance, virulence, thermal tolerance and osmotic stress tolerance (17). Thus, C. *auris* might have previously existed as a plant saprophyte in specialized ecosystems such as the wet lands. As a first step, its emergence might have been linked to global warming (including climatic oscillations) effects on wetlands (18), and its enrichment in that ecological niche was the result of *C. auris* combined thermal tolerance and salinity tolerance. Interestingly, areas where *C. auris* was first recognized overlap, at least in part, with the impacted wet land ecosystems (18). Of interest, *C. albicans* can be a part of wet land ecosystem (19), and although many of the virulence determinants in the *C. auris* genome have not been characterized, it is theoretically possible that human pathogenic *Candida* spp. have passed some virulence traits to previously non-pathogenic *C. auris* through plasmid DNA transfer (20,21), in the backdrop of changing ecological niches. Alternatively, the effect of higher ultraviolet radiation in combination with global warming (22), might have contributed to mutagenic events that resulted in suddenly increased fitness of a saprobe for survival in host, via melanin or non-melanin dependent processes (23). *C. auris* jump from an environmental fungus to transmission and pathogenicity in humans might have had an intermediate host, specifically an avian host as fungi that can grow in 40 or 42 ° C can infect avian fauna. Of note, sea birds may serve as reservoirs for indirect transmission of drug resistant *Candida* species, such as *C. glabrata*, to humans (24). The uncanny ability of C *auris* for niche-specific adaptations, first in the environment, then in an avian host might have led as a third step to the ultimate establishment as a human pathogens through genetic and epigenetic switches (25). The hypothesis that *C. auris* broke the mammalian thermal barrier through adaptation to climate change suggests several experimental lines of investigation to obtain evidence for and against it. If *C. auris* thermal tolerance is indeed a new property, a careful analysis of the temperature range where it grows might reveal that it is still less tolerant than *Candida* spp. that have a long association with mammals. In this regard, isolates from earlier outbreaks may show differences in thermal tolerance from those of outbreaks that are more recent. Of course, analysis of thermal susceptibility would require a more detailed experimental approach to identify small differences than are normally measured by routine microbiologic characterization. A search for environmental reservoirs could produce closely related *C. auris* strains that have not yet adapted to higher temperatures, which may be expected if the process is stochastic and only some clades have made the transition. Increased sampling in these environments and hosts respectively could test involvement of wetlands and/or birds. Should evidence be found for *C. auris* or close relatives in these environments, then one might consider that the host jump from birds to humans follows similar mechanisms to those that operate for influenza virus. The possibility that *C. auris* was always a health care pathogen that has been recently discovered appears to have little support given that no isolate was found prior to 1996 in fungal collections (3). Evaluating the contributory role of high human population densities, migrations, increased city temperatures (15), low hygiene, pollution, regional and international travel (26) in the emergence of *C. auris* is difficult, given the available information, but these potential associations are fertile areas for research. The emergence of *C. auris* also poses interesting basic science questions on the thermal stability of its enzymatic processes, and in a broader sense, the mechanisms of virulence and adaptation for human pathogenic fungi. For example, the evolutionarily conserved Hog1 stress-activated protein kinase (SAPK) whose role in *C. neoformans* virulence is established (27), promotes stress resistance and virulence in *C. auris* (28), suggesting tantalizing connections between mechanisms for adaptation and virulence that are ripe for further study. A scheme for the possible factors operating on the emergence of *C. auris* is shown in Figure 2.

**Figure 2.**
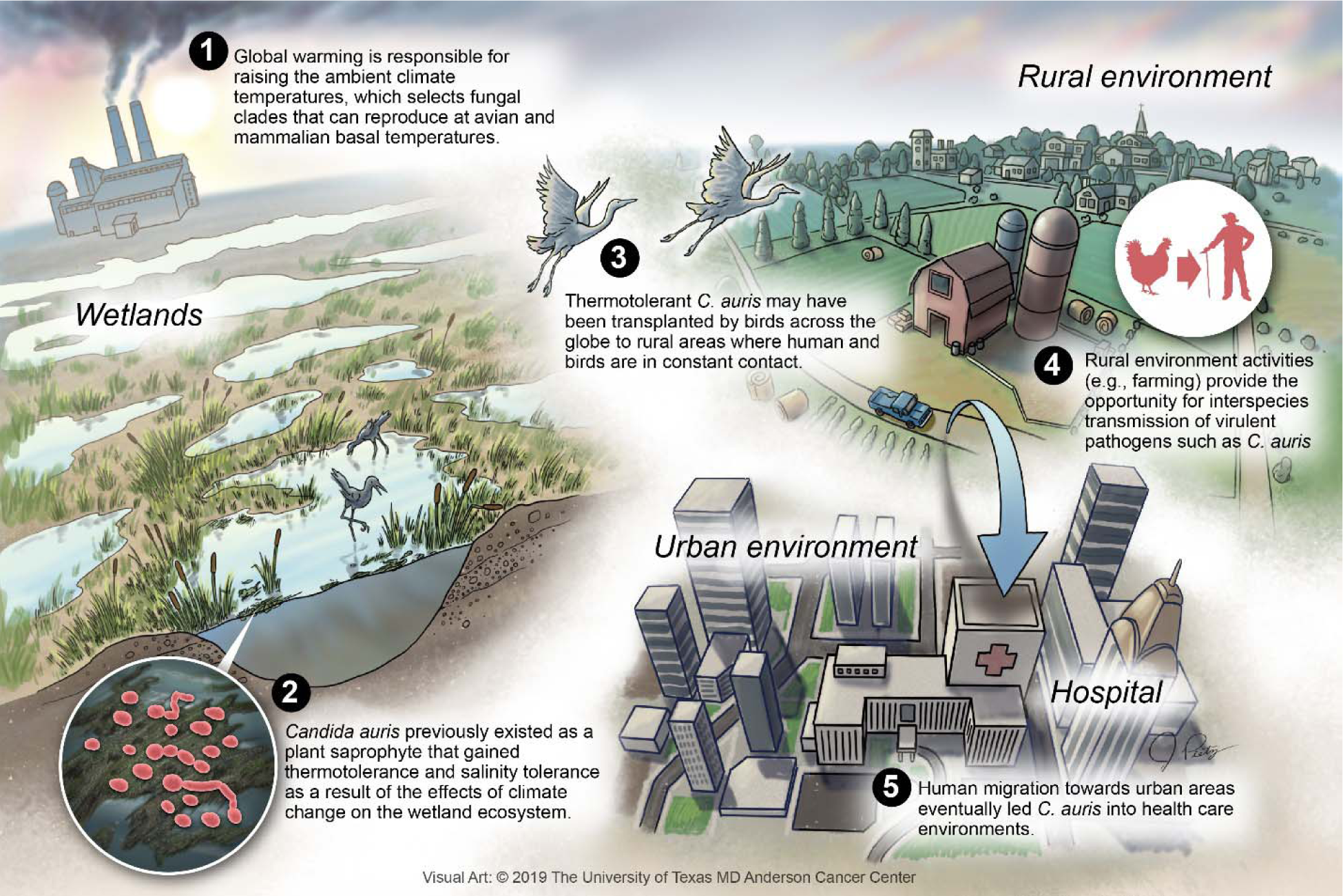
Proposed scheme for the emergence of *C. auris*.

Although global warming-related changes in the environment might have played a prominent role in *C. auris* emergence, this variable is unlikely to explain the whole story. For example, it is difficult to see how global warming alone explains the spontaneous emergence of four clades of *C. auris* in geographically disparate regions, each separated by thousands of years of evolutionary distance to each other, unless there is another common epidemiological variable that facilitated the interaction with humans for virulence to become apparent. Finally, we note that all four clades contain MTLa and MTLa loci (29), which raise the possibility that they could interact through mating if their current geographic isolation is ended by inadvertent human transport or migratory bird patterns.

Whether *C. auris* is the first example of new pathogenic fungi emerging from climate change or whether its origin into the realm of human pathogenic fungi followed a different trajectory, its emanation stokes worries that humanity could be facing new diseases from fungal adaptation to hotter climates. Thus far, the majority of human cases of *C. auris*-related disease have occurred in debilitated individuals such as those in intensive care units. Because their debilitated condition impairs their immunity, this group could serve as sentinels for the appearance of new fungal diseases. Perhaps the greatest lesson from the emergence of *C. auris* is the need for greater vigilance and continuous monitoring. In this regard, the environment is likely to contain large numbers of fungal species with pathogenic potential that are currently non-pathogenic for humans because they lack the ability to grow at mammalian temperatures. If anything, the direct and indirect effects of climate changes induced by an exponentially growing human population as drivers of fungal evolution should be an area of intense research in the decades to come. Widening of geographic range of both innately thermal tolerant pathogenic fungi and the acquisition of virulence traits in thermal tolerant non-pathogenic environmental fungi could shape the 21^st^ century as an era of expanding fungal disease for both the fauna and flora of the planet.

